# Adversarial Learning For End-To-End Cochlear Speech Denoising Using Lightweight Deep Learning Models

**DOI:** 10.1101/2025.03.19.644108

**Authors:** Tom Gajecki, Waldo Nogueira

## Abstract

This paper investigates an end-to-end speech signal denoising approach for cochlear implants (CIs). Building on previous work, we first explore the effect of relocating the deep envelope detector within the deep learning-based CI sound coding strategy, moving it from the skip connection to the output of the masking operation. This modification enables high-resolution time-frequency masking and optimizes noise reduction. Next, we introduce a discriminator network to further enhance the model by enforcing the generation of higher-quality electrodograms (i.e., the electric pulse patterns that stimulate the auditory nerve). This adversarial learning approach improves the generation of electrodograms and has the potential to enhance speech understanding for CI users. Objective evaluations, including signal-to-noise ratio improvement and linear cross-correlation coefficients, demonstrate that these enhancements significantly boost the performance of the end-to-end CI speech-denoising algorithm while reducing its parameter count, making it suitable for real-time applications.

## 1. INTRODUCTION

Cochlear implants (CIs) have long been a solution for individuals with profound sensorineural hearing loss, enabling them to regain the sensation of hearing through direct electrical stimulation of the auditory nerve [1]. Despite significant advancements in CI technology, a persistent challenge remains in noisy environments, where traditional sound coding strategies, such as the Advanced Combination Encoder (ACE), often struggle to maintain speech intelligibility. Recent approaches utilizing deep learning for end-to-end speech denoising in CIs have shown great potential in enhancing noise reduction by generating high-quality electrodograms directly from raw audio signals [2, 3]. However, many of these models are computationally intensive, which can hinder their real-time applicability.

In this paper, we present an extension of an end-to-end CI speech denoising sound coding strategy [4]. While the end-to-end CI processing explored here can be applied to any commercial CI sound coding strategy, our focus is on ACE (i.e. Deep ACE). We begin by examining the impact of relocating the deep envelope detector (DED) from the skip connection to the output of the masking operation. Next, we introduce an adversarial version of our end-to-end speech denoising system, following a similar approaches such as [5, 6]. This new model contains a discriminator and acts as an adversarial network, helping to ensure that the electrodograms are both noise-reduced and better aligned with the underlying speech signals.

Furthermore, we provide a theoretical analysis suggesting that electrodograms are inherently less complex than raw audio signals. This reduction in complexity makes electrodograms easier for neural networks to predict, as the network can focus on capturing the most relevant features for speech intelligibility without being overwhelmed by the intricacies of raw audio. By simplifying the prediction task, the network can allocate more resources to refining the electrodogram quality, thereby improving its ability to preserve crucial speech information.

The proposed model is evaluated through a series of objective measures, demonstrating that the adversarial deep learning-based CI speech denoising model outperforms the used baselines. This approach offers a more efficient solution for potential real-time deployment, with improvements observed in noise reduction and the quality of generated electrodograms. Although listening tests were not conducted, the objective results suggest that our model could significantly enhance speech intelligibility and overall user experience in complex auditory environments.

## 2. METHODS & MATERIALS

### 2.1. Algorithms

#### Deep ACE V2

This is an enhanced version of the original Deep ACE model [4] (see Figure 1a), designed to improve noise reduction and electrodogram quality for CIs. A key innovation in Deep ACE V2 is the repositioning of the Deep Envelope Detector (DED; Figure 1b) to occur after the masking operation. This adjustment allows the model to leverage high-resolution time-frequency masking, capturing more detailed speech features and filtering out noise more effectively. By placing the DED after the masking stage, Deep ACE V2 generates electrodograms that are more refined and closely aligned with the desired outputs, leading to better speech intelligibility in noisy environments. This configuration also lays the foundation for further enhancements, such as the integration of a discriminator network, as explored in this study.

**Fig. 1.**
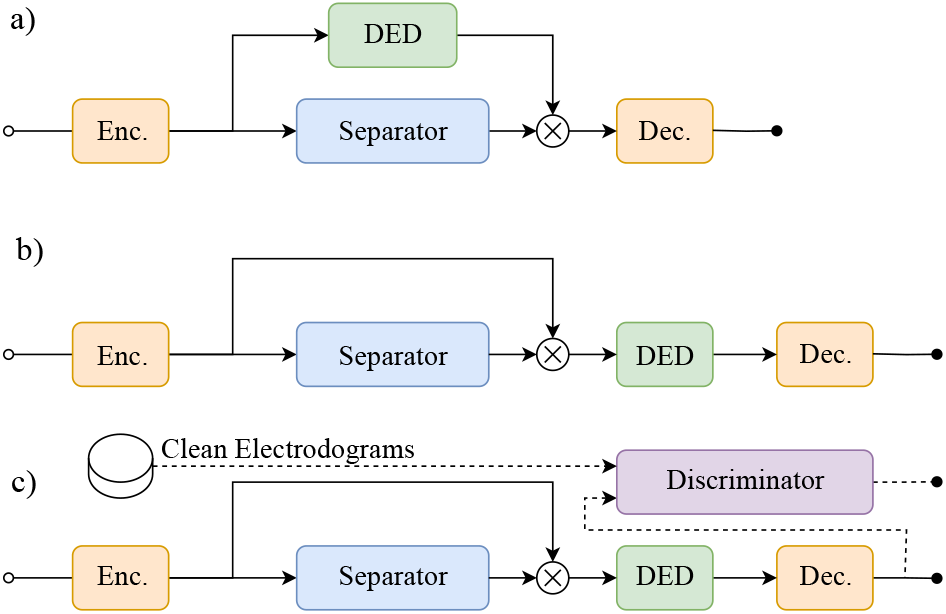
Deep learning modules of the investigated end-to-end CI speech denoising variants. The original Deep ACE with the deep envelope detector (DED) in the skip connection (a), the here presented Deep ACE V2 with the DED after the masking operation (b), and the Adversarial Deep ACE (c).

#### Adversarial Deep ACE model

In this model, a multi-scale convolutional discriminator network (inspired by [7]) is introduced to enhance electrodogram generation (see Figure 1c). Acting as an adversarial network, the discriminator distinguishes between realistic and unrealistic electrodograms, guiding the generator to produce higher-quality outputs that better mimic auditory signals. The discriminator operates at multiple resolutions, using downsampling to capture both fine and coarse details while applying normalization techniques to ensure stable training. Notably, the discriminator is only used during training, maintaining a low computational footprint for real-time inference. Combined with the strategic placement of the DED after the masking operation, this approach enables Adversarial Deep ACE to leverage high-resolution time-frequency masking, resulting in improved noise reduction and enhanced speech intelligibility in noisy environments.

### 2.2 Loss Functions for adversarial Speech Denoising

The training of the Adversarial Deep ACE model involves several key loss functions that collectively contribute to generating high-quality electrodograms for CIs. The training configuration for the other models used in this study is consistent with the procedures described in [4].

#### Adversarial Loss (ℒ _adv_)

The adversarial loss is crucial for training the generator to produce output signals that are indistinguishable from real ones by the discriminator. It is calculated using binary cross-entropy (BCE), averaged over all discriminators. Specifically, the adversarial loss is defined as:

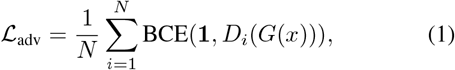

where *N* represents the number of discriminators, *D*_*i*_(*G*(*x*)_*i*_) denotes the discriminator’s output for the generated denoised speech from the noisy audio signal *x* at the *i*-th discriminator, and **1** is a vector of ones, representing the target label for real target data.

#### Reconstruction Loss (ℒ _recon_)

The reconstruction loss measures the discrepancy between the target signals and those generated by the model. In Adversarial Deep ACE, this loss is computed using the Mean Squared Error (MSE) between the generated *G*(*x*) and the target signals. This ensures that the generated signals closely align with the target signals, improving the quality of the output electrodograms. In contrast, Adversarial TasNet uses the Scale-Invariant Signal-to-Distortion Ratio (SI-SDR; [8]) as its reconstruction loss, optimizing the generated signals to better match the clean target audio.

#### Feature Matching Loss (ℒ _feat match_)

Feature matching loss [9] is introduced to encourage the generator to produce electrodograms that match the feature space of real electrodograms. This is calculated as:

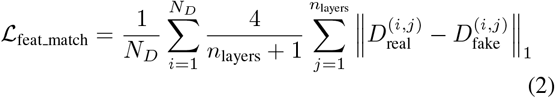

Here, *N*_*D*_ is the number of discriminators, and *n*_layers_ is the number of layers within each discriminator. The outputs 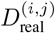 and 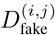 represent the discriminator’s response to real and generated electrodograms at the *i*-th discriminator and *j*-th layer, respectively. The *L*_1_ norm is used to measure the absolute difference between these features, ensuring that the generated electrodograms not only look realistic but also possess similar internal feature structures as real electrodograms.

#### Total Loss

The total loss combines adversarial, reconstruction, and feature-matching losses, weighted empirically as 10.0 for feature matching and 1.0 for the others, which balance contributions to ensure high-quality outputs.

The models in this work are designed to be lightweight, as summarized in Table 1. For a proper baseline comparison with Adversarial Deep ACE, we employ an adversarial version of TasNet, which uses the same discriminator as the one in Adversarial Deep ACE and is trained similarly. The table provides the number of parameters for both the generator and, where applicable, the discriminator. It is important to highlight that the discriminator is only utilized during training to improve the quality of the generated electrodograms, but it is not required during inference. This contributes to the system’s efficiency. By maintaining a low parameter count, these models strike a balance between computational efficiency and performance, making them well-suited for real-time applications.

**Table 1.**
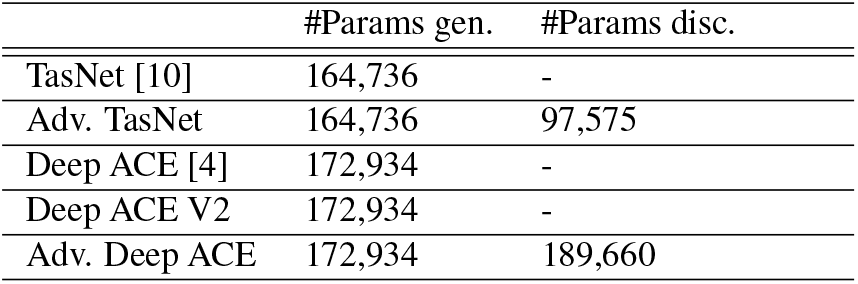
Comparison of Generator (gen.) and Discriminator (disc.) parameter counts for various models This table presents the number of parameters for both the generator and discriminator across different models.

|  | #Params gen. | #Params disc. |
| --- | --- | --- |
| TasNet [10] | 164,736 | - |
| Adv. TasNet | 164,736 | 97,575 |
| Deep ACE [4] | 172,934 | - |
| Deep ACE V2 | 172,934 | - |
| Adv. Deep ACE | 172,934 | 189,660 |

### 2.3. Objective Instrumental Measures

***SNRi***: To evaluate the noise reduction achieved by each of the tested algorithms, we compute the SNR improvement (SNRi). This metric is calculated in the electrodogram domain and compares the original input SNR with the SNR obtained after denoising. The SNRi is defined as:

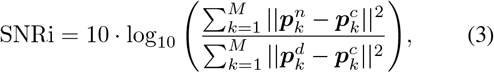

where ***p****k* represents the LGF output of band *k*, and the superscripts *n, c*, and *d* correspond to the noisy, clean, and denoised electrodograms, respectively.

Similarly, SNRi can also be computed in the audio domain by replacing the LGF output with the predicted audio signals. This involves comparing the SNR of the original noisy audio signal to the SNR of the denoised audio output, using the predicted audio in place of electrodograms.

#### Linear Cross Correlation (LCC)

To evaluate distortions and artifacts from the tested algorithms, we calculated the linear correlation coefficients (LCCs) between the clean ACE electrodograms (***p***^*c*^) and the denoised electrodograms (***p***^*d*^). The LCCs were computed for each of the 22 channels to assess degradation in individual channel outputs caused by the denoising process. The LCC for channel *k* (LCC_*k*_) is calculated using the Pearson correlation coefficient [11] as follows:

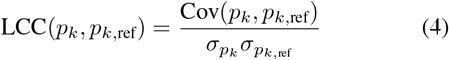

where Cov(*p*_*k*_, *p*_*k*,ref_) is the covariance between the predicted and reference electrodograms at frame *k*, and *σ*_*p*_*k* and *σ*_*p*_*k*,ref are their standard deviations. A higher LCC indicates a stronger linear relationship, reflecting the predicted electrodograms’ accuracy in capturing the reference signals’ underlying patterns. For the TasNet variants, the LGF output is replaced by the estimated denoised speech audio signal.

#### Complexity Measures

To assess the complexity of the predicted electrodograms and audio samples, we employ two complementary measures: *Lempel-Ziv complexity* and *Shannon entropy*. Each measure provides a different perspective on the variability and unpredictability of the sequences.

*Lempel-Ziv complexity (LZC)* quantifies the complexity of the predicted electrodogram sequences and audio samples by counting the number of distinct patterns or configurations observed throughout the sequence. For electrodograms, LZC measures the diversity of electrode activation patterns, while for audio signals, it evaluates the occurrence of new, previously unseen patterns in the time-domain waveform. A higher LZC value indicates a more complex and less predictable sequence, reflecting greater variability and richness in both the electrodograms and audio signals.

*Shannon entropy* measures the unpredictability or information content in the predicted electrodogram sequences and audio samples. Entropy is calculated as:

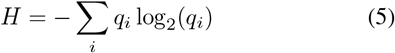

where *q*_*i*_ represents the probability of occurrence for each unique pattern. In electrodograms, entropy assesses the distribution of electrode activation patterns across frames, while in audio samples, it evaluates the variability in signal amplitude or frequency content. Higher entropy values suggest more complex and less predictable sequences, indicating greater variability in both electrodograms and audio signals.

### 2.4. Audio Data

In this study, we trained the models using speech signals from the LibriVox corpus [12] and mixed them with noise from the DEMAND dataset [13]. For testing, we utilized the Hochmair, Schulz, Moser (HSM) sentence dataset, which was mixed with CCIT speech-shaped noise [14] and ICRA7 babble noise [15]. The noise levels were adjusted to a signal-to-noise ratio (SNR) range of -5 to 10 dB.

## 3. RESULTS

### SNRi

Figure 2 presents the SNRi achieved by the evaluated algorithms. The results show that the proposed Adversarial Deep ACE method consistently outperforms the baseline algorithms across all input SNRs for both CCITT and ICRA7 noise conditions. Interestingly, the adversarial TasNet performs worse than the standard TasNet in CCITT noise but performs comparably in ICRA7. This discrepancy may be attributed to suboptimal tuning of the adversarial model for specific noise types, indicating the need for further optimization or task-specific adjustments.

**Fig. 2.**
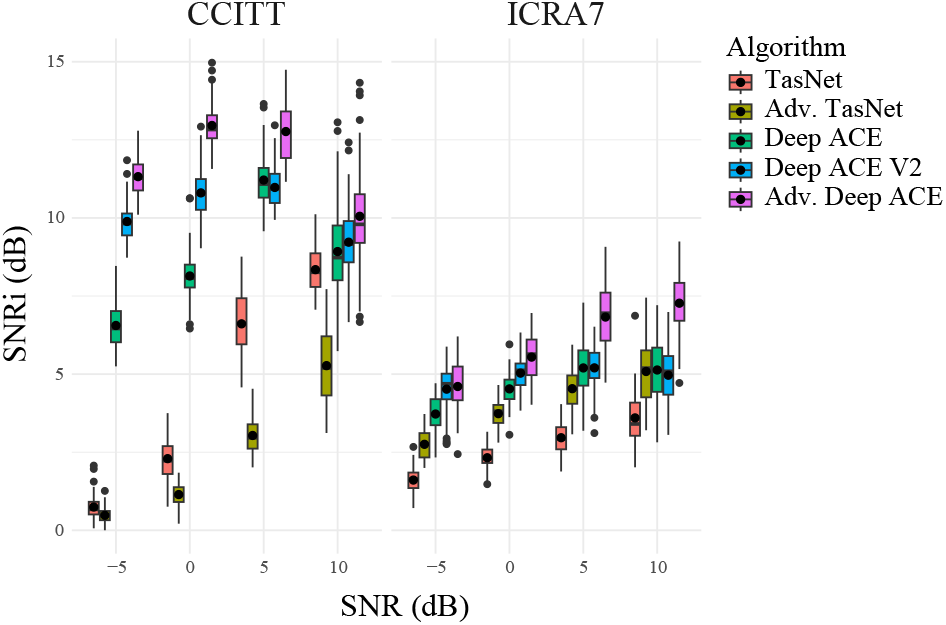
Box plots showing the SNRi scores in dB for the tested algorithms in SSN and ICRA7 noises for the different SNRs using the HSM speech dataset.

#### LCCs

Figure 3 shows the LCCs as a function of electrode number for each algorithm. The results indicate that CI denoising strategies perform best at lower frequencies, likely due to their focus on enhancing low-frequency modulations, which are critical for speech understanding [16]. In contrast, TasNet shows a slight advantage at higher frequencies, which are important for capturing consonants and environmental sounds.

**Fig. 3.**
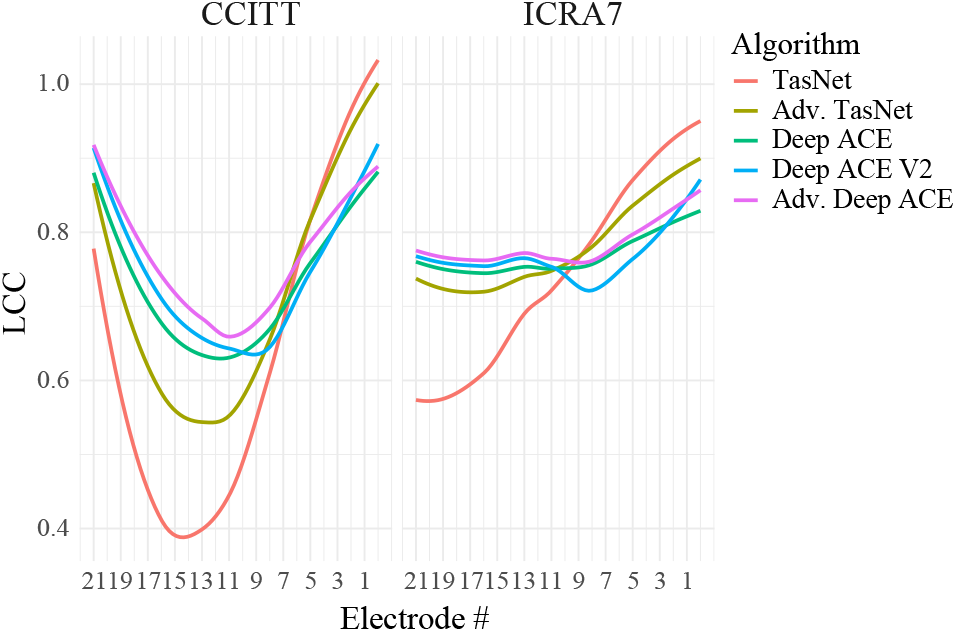
Polynomial regressions showing the channel-wise LCCs between processed and clean electrodograms for the different algorithms and noises using the HSM dataset. Higher electrode numbers correspond to higher frequencies.

#### Complexity measures

Figure 4 shows violin plots comparing the complexity of audio signals and electrodograms. The complexity measures, based on LZC and Shannon entropy, reveal that electrodograms have lower complexity than audio signals. This simplification likely explains the superior performance of CI denoising strategies, which focus on predicting electrodograms. In contrast, TasNet, which directly predicts audio, must handle higher complexity, making denoising more challenging in noisy environments. Predicting electrodograms offers a more manageable signal, potentially improving CI denoising effectiveness.

**Fig. 4.**
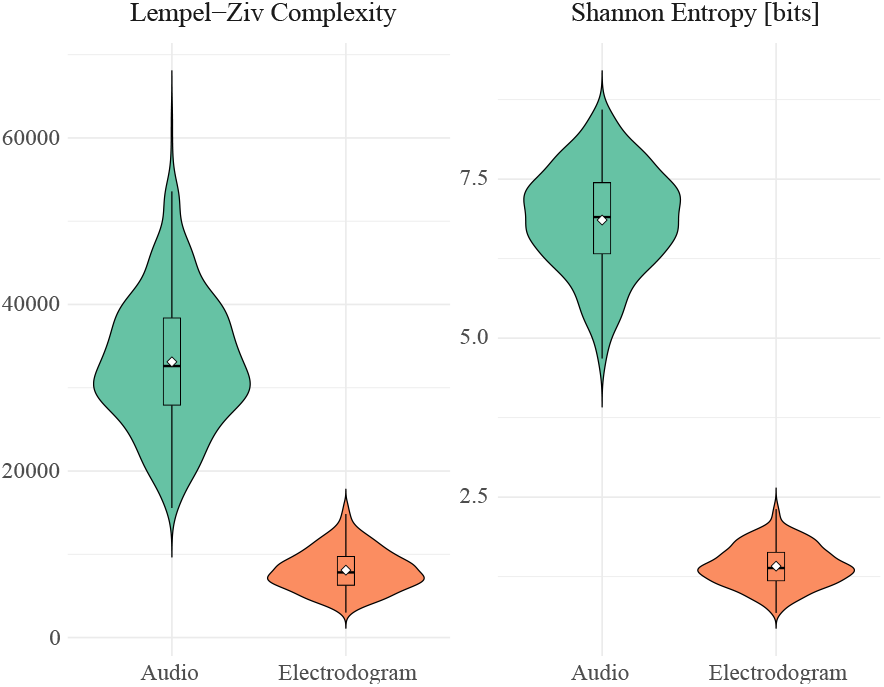
Violin plots comparing the complexity of audio signals and electrodograms using Lempel-Ziv complexity (LZC) and Shannon entropy.

## 4. CONCLUSIONS

In this study, we enhanced a deep learning-based denoising CI sound coding strategy by integrating a discriminator network and repositioning the DED, improving noise reduction and electrodogram quality. Objective measures like SNR improvement and LCCs show that the adversarial CI sound coding strategy outperforms the used baselines. The simplicity of electrodograms compared to raw audio likely contributes to this improved performance. These advancements represent a promising step toward more effective, real-time speech enhancement in noisy environments. Future work could focus on the optimization and validation through listening tests.

